# Fending for oneself or partnering up: Competition between mixo- and heterotrophic ciliates under dynamic resource supply

**DOI:** 10.1101/2023.11.06.564392

**Authors:** Sabine Flöder, Toni Klauschies, Moritz Klaassen, Tjardo Stoffers, Max Lambrecht, Stefanie Moorthi

## Abstract

The outcome of species competition strongly depends on the traits of the competitors and associated trade-offs, as well as on environmental variability. Here we investigate the relevance of consumer trait variation for species coexistence in a ciliate consumer – microalgal prey system under fluctuating regimes of resource supply. We focus on consumer competition and feeding traits, and specifically on the consumer’s ability to overcome periods of resource limitation by mixotrophy, i.e. the ability of photosynthetic carbon fixation via algal symbionts in addition to phagotrophy. In a 48-day chemostat experiment, we investigated competitive interactions of different heterotrophic and mixotrophic ciliates of the genera *Euplotes* and *Coleps* under different resource regimes, providing prey either continuously or in pulses under constant or fluctuating light, entailing periods of resource depletion in fluctuating environments, but overall providing the same amount of prey and light. Although ultimate competition results remained unaffected, population dynamics of mixotrophic and heterotrophic ciliates were significantly altered by resource supply mode. However, the effects differed among species combinations and changed over time. Whether mixotrophs or heterotrophs dominated in competition strongly depended on the genera of the competing species and thus species-specific differences in the minimum resource requirements that are associated with feeding on shared prey, nutrient uptake, light harvesting and access to additional resources such as bacteria. Potential differences in the curvature of the species’ resource-dependent growth functions may have further mediated the species-specific responses to the different resource supply modes. In addition, while the presence of a heterotrophic competitor may have a direct negative effect on the growth rate of a mixotrophic species through grazing on a shared prey species, its presence may also have an indirect positive effect on the growth rate of the mixotroph by reducing competition between the autotroph and mixotroph for shared nutrients and light. Our study thus demonstrates that complex trophic interactions determine the outcome of competition, which can only be understood by taking on a multidimensional trait perspective.

## INTRODUCTION

Mixotrophy is pervasive and common in both aquatic and terrestrial ecosystems (Selosse et al. 2017). Combining two functionally different modes of nutrition, i.e., phototrophy and heterotrophy, mixotrophic organisms are able to use inorganic and organic carbon sources for biomass production (Esteban et al. 2010, Andersen et al. 2015). Although research on organisms characterized by this combination of nutritional strategies has a long history, results of more recent studies suggest that the contribution of mixotrophs to trophic interactions, ecosystem functioning and biogeochemical cycles is much higher than previously assumed (e.g. Caron et al. 2012, Flynn et al. 2013, Mitra et al. 2014, Stoecker et al. 2017).

Combining phagotrophy and phototrophy within a single cell, mixotrophic protists contribute largely to aquatic primary production (Jansson et al. 1996, Laybourn-Parry et al. 1997, Burkholder et al. 2008, Wilken et al. 2013) and exert grazing pressure on the microbial community (e.g. Zubkov and Tarran 2008, Mitra et al. 2014, Unrein et al. 2014). The degree of mixotrophy varies among species and has been described as a gradient ranging from primarily phototrophic to primarily heterotrophic (e.g. Sanders et al. 1990, Jones 1994, Flynn et al. 2013). Within species, the relative importance of primary production and phagotrophy is affected by prey abundance, light intensity and macronutrient concentration (Sanders et al. 1990, Caron et al. 1993, Jones 1994, Urabe et al. 2000, Smalley et al. 2003). The tremendous functional diversity of mixotrophs that exists today results from the fact that the ability to photosynthesise has been acquired, lost and reacquired several times during the evolutionary development of protist groups (Archibald 2009, Figueroa-Martinez et al. 2015, Leles et al. 2018, Flynn et al. 2019, Hampl et al. 2019). Therefore, mixotrophic protists have been recently grouped into constitutive mixotrophs (CM) with an innate capacity for photosynthesis, and different categories of non-constitutive mixotrophs (NCM) that acquire their photosystems in different ways (Mitra et al. 2016).

Supplementing or substituting photosynthetic energy supply under limiting light conditions, and heterotrophic energy supply during prey shortages, mixotrophic protists gain starvation resistance, which facilitates growth or survival under resource depletion (e.g. Bird and Kalf 1987, Smalley et al 2003, Flöder et al 2006). Accordingly, mixotrophy should be the superior nutritional strategy under fluctuating environmental conditions where the resource supply varies over time. In contrast, mixotrophs have been considered inferior to their purely heterotrophic and photoautotrophic counterparts based on the assumption that pursuing two different nutritional strategies infers metabolic investments into different nutritional traits that are limited by total cell resource and thus result in energetic trade-offs (e.g. Dolan and Pérez 2000, Calbet et al. 2011, Andersen et al. 2015). As a consequence, heterotrophs and photoautotrophs can be expected to be superior under constant environmental conditions comprising sufficient supply of prey, light and nutrients, respectively. Despite the broad range of functional diversity among mixotrophic protists, the assumption of an energetic trade-off is widely supported.

Most evidence for the energetic trade-off assumption comes from comparative studies on CM flagellates. The CM *Poterioochromonas malhamensis* displayed lower maximum growth rates and slightly lower ingestion rates than the heterotrophic flagellate *Spumella elongata* irrespective of light conditions, however accumulating more biomass than the heterotroph when light was available (Pålsson and Daniel 2004). Regarding photosynthesis, different CMs exhibited different capabilities to grow without prey; however, oxygen production per unit biovolume was low compared to pure photoautotrophs, indicating a lower photosynthetic capacity (Rottberger et al. 2013). In competition experiments between the heterotrophic flagellate *Spumella* and the CM flagellate *Ochromonas*, light availability determined their competitive outcome. When only phagotrophic growth was possible, the heterotroph excluded the mixotroph, while the opposite was observed under a high light : bacteria supply ratio. Simultaneous supply of both resources resulted in coexistence, but a higher biomass of the CM due to its supplementary photosynthetic capabilities. Competing with the pure phototroph *Cryptomonas* and bacteria for dissolved phosphorus (P), *Ochromonas* was able to coexist by feeding on the P-rich bacteria, taking up P but at the same time eliminating another competitor for P (bacteria) (Rothhaupt 1996). These studies indicate that performance and competition of mixotrophs is determined by the amount of alternative resources the mixotroph is capable of using, e. g., light and dissolved inorganic nutrients in competition with heterotrophs, and prey organisms in competition with phototrophs. Supporting this observation, previous studies have indicated that light, which determines photosynthetic carbon fixation, mediates the impact of mixotrophs by altering their requirements for organic carbon sources such as prey (Tittel et al. 2003, Ptacnik et al. 2016) and determines their success in natural plankton communities (Edwards 2019).

Rather than investing into two metabolic pathways of their own, NCMs obtain their photosystems from different sources, i.e. by either retaining chloroplasts from specific or several different prey organisms (plastidic specialist non-constitutive mixotroph, pSNCM and generalist non-constitutive mixotroph, GNCM, respectively) (Mitra et al. 2016). Without the need to synthesize the photosynthetic apparatus themselves, energetic costs in acquired phototrophs are likely to be reduced, and the costs of chloroplast retention can be adjusted by slowing plastid turnover when food supply is low (Mcmanus et al. 2012). However, obligate ciliate GNCMs and pure heterotrophic marine ciliates were shown to exhibit similar overall growth rates despite the energy subsidy received by mixotrophs via photosynthesis from retained chloroplasts (Schoener and McManus 2017). The authors suggested that this might be due to the fact that obligate mixotrophic growth is not possible in the dark under light : dark diel cycles. Therefore, benefits though mixotrophic nutrition that can only be gained during the light phase are not sufficient to exceed potential daily costs of symbiont maintenance and respiration. They postulated that the advantage of chloroplast retention is only high when food concentrations are low, but diminishes when food concentrations are high. Under food limitation, GNCMs were shown to be more resistant than their heterotrophic counterparts (McManus et al. 2018, Maselli et al. 2020, Hughes et al. 2021). In contrast, GNCMs are generally competitively inferior under food replenished conditions, supporting the assumption of an energetic trade-off.

Protists hosting photoautotrophic endosymbionts (eSNCM) (Mitra et al. 2016) take an exceptional position among mixotrophs, as phagotrophic and phototrophic traits are provided by individual organisms. Trophic associations of ciliates and endosymbiontic algae provide a close nutritional coupling, with inorganic nutrients passing from the host to the symbionts and photosynthesis products from the symbionts to the host (Reisser 1992). Freshwater ciliates predominantly host green endosymbionts including different genera like *Choricystis*, *Micractinium* and *Chlorella* (Hoshina et al. 2005, Pröschold et al. 2011). Comparisons of mixotrophs and heterotrophs have largely been based on different species with genus- or species-specific traits potentially confounding differences in performance based on heterotrophic or mixotrophic nutrition. Endosymbiotic mixotrophs enable the comparison of symbiont-bearing and symbiont-free organisms of the same genus, or even the same species. The symbiont-bearing ciliate *P. bursaria*, for instance, exhibited lower ingestion rates than symbiont-free populations of the same species (Berk et al. 1991, Miura et al. 2017), which, however, depended on light intensity, approaching levels similar to those of the heterotrophic *P. bursaria* at high light (Berk et al. 1991). Receiving metabolic benefits from their symbionts has been shown to increase the competitive ability of the eSNCM *P. bursaria* and to facilitate its coexistence with the heterotroph *Colpidium* sp. via niche expansion (Hsu and Moeller 2021, Hsu et al. 2022). Again, light availability determined competition between heterotrophs and symbiont-bearing mixotrophs, promoting the mixotroph at high light (Hsu et al. 2022), while photosynthetic carbon fixation prolonged its survival during periods of prey depletion (Iwai et al. 2019). Taken together, the type and the quantity of resource supply seems to determine mixotroph performance and competitive ability. Most studies, however, have been performed under constant environmental conditions, although resource supply is often highly variable in nature, so that adaptive strategies to overcome periods of resource limitation can be beneficial in variable environments (Flöder et al. 2018).

In the current study, we therefore investigated population dynamics of competing pairs of heterotrophic and algal endosymbiont bearing ciliates in a 48-day chemostat experiment under constant and fluctuating light and prey supply, using the heterotrophic *Euplotes octocarinatus* and *Coleps hirtus*, and the eSNCM *Euplotes daidaleos* and *Coleps hirtus viridis*. Incubating two same-genus and two mixed-genus combinations of heterotrophs and eSNCMs to the different resource supply regimes, we aimed at elucidating how the outcome of competition depends on the overall trait similarity among competitors. We expected the outcome of competition among heterotrophic and symbiont-bearing ciliates to depend 1) on the mode of resource supply (constant versus fluctuating / pulsed), and 2) on the specific ciliate combination, i.e. their specific traits as well as their similarity (same-genus, mixed-genus combination). Specifically, we expected pulsed prey supply entailing periods of resource depletion to promote the mixotrophs based on their starvation resistance due to photosynthetic carbon fixation, especially at constant light supply and under periods of high light supply. We further expected resource fluctuations (both prey and light) to promote coexistence of both species. Regarding species composition, we assumed the outcome of competition for ciliates of the same genus to primarily depend on the resource supply regime, while ciliates of different genera might be less affected by resource supply but rather by other functional traits characteristic to the different genera.

## MATERIALS AND METHODS

### Organisms used and culture conditions

We used four freshwater ciliates belonging to the genera *Coleps* and *Euplotes* that differed in trophic mode, feeding preference and average cell size (see Table 1 for characteristics and origin of the organisms used). *Coleps hirtus* and *Euplotes octocarinatus* are heterotrophic, whereas *Coleps viridis* and *Euplotes daidaleos* are mixotrophic. Both ciliates carry phototrophic symbionts. We did not perform sequence analysis on the symbionts of our *E. daidaleos* and *Coleps viridis* strains. However, *Chlorella vulgaris* has been identified for other *E. daidaleos* strains, whereas *Micractinium conductrix* is the endosymbiont of several *Coleps viridis* strains (Pröschold et al. 2011, Pröschold et al. 2021). None of the cultures used are axenic and are accompanied by a microbial community consisting of bacteria and heterotrophic flagellates. All ciliates were fed with the microalga *Cryptomonas* sp. (also not axenic). Both of the *Euplotes* and the *Coleps* species used in this experiment are also able feed on bacteria; however, only *Euplotes* is able to grow on a diet consisting solely of bacteria. Abiotic resources were light and mineral nutrients. Cultures of *Cry* were grown in WEES medium (Kies 1967), mineral water (Volvic) served as culture medium for ciliates.

**Table 1:** Three-factorial linear mixed model ANOVA results. Response variable: log-ratio mixotrophic to heterotrophic biovolume. Species combination (C), and supply modes of light (L) and prey (P) were factors, time (T) was set as trend and as factor to account for random fluctuations.

**Type III Analysis of Variance Table**
|  | Sum Sq | Mean Sq | Num-DF | Den-DF | F-value | P-value |
| --- | --- | --- | --- | --- | --- | --- |
| **C** | 27.49 | 9.17 | 3 | 81.26 | 42.1 | <2.2e<sup>-16</sup> |
| **L** | 0.19 | 0.187 | 1 | 81.24 | 0.856 | 0.358 |
| **P** | 0.91 | 0.910 | 1 | 81.23 | 4.18 | 0.044 |
| **T** | 0.30 | 0.296 | 1 | 23.62 | 1.36 | 0.255 |
| **C x L** | 2.27 | 0.755 | 3 | 81.26 | 3.47 | 0.020 |
| **C x P** | 0.62 | 0.208 | 3 | 81.25 | 0.955 | 0.418 |
| **L x P** | 0.20 | 0.195 | 1 | 81.23 | 0.897 | 0.346 |
| **C x T** | 439.61 | 147 | 3 | 1054 | 673 | <2.2e<sup>-16</sup> |
| **L x T** | 0.03 | 0.031 | 1 | 1055 | 0.143 | 0.706 |
| **P x T** | 1.71 | 1.72 | 1 | 1054 | 7.88 | 0.005 |
| **C x L x P** | 0.29 | 0.095 | 3 | 81.3 | 0.438 | 0.726 |
| **C x L x T** | 3.62 | 1.21 | 3 | 1054 | 5.54 | 0.001 |
| **C x P x T** | 6.80 | 2.27 | 3 | 1054 | 10.4 | 9.352e<sup>-07</sup> |
| **L x P x T** | 4.44 | 4.44 | 1 | 1054 | 20.4 | 7.009e<sup>-06</sup> |
| **C x L x P x T** | 3.34 | 1.12 | 3 | 1054 | 5.11 | 0.002 |

**Contrasts**
|  | estimate | SE | df | t-ratio | p-value |
| --- | --- | --- | --- | --- | --- |
| **L = continuous, P = constant** |  |  |  |  |  |
| C. mixo – C. het <> C. mixo – E. het | 0.3152 | 0.154 | 29.7 | 2.040 | 0.1965 |
| C. mixo – C. het <> E. mixo – E. het | -1.1533 | 0.154 | 29.7 | -7.465 | <.0001 |
| C. mixo – C. het <> E. mixo – C. het | -2.1020 | 0.154 | 29.7 | -13.61 | <.0001 |
| C. mixo – E. het <> E. mixo – E. het | -1.4684 | 0.154 | 29.7 | -9.505 | <.0001 |
| C. mixo – E. het <> E. mixo – C. het | -2.4171 | 0.154 | 29.7 | -15.65 | <.0001 |
| E. mixo – E. het <> E. mixo – C. het | -0.9487 | 0.154 | 29.7 | -6.141 | <.0001 |
| **L = fluctuating, P = constant** |  |  |  |  |  |
| C. mixo – C. het <> C. mixo – E. het | 0.0454 | 0.174 | 30.6 | 0.261 | 0.9937 |
| C. mixo – C. het <> E. mixo – E. het | -1.3177 | 0.173 | 29.7 | -7.629 | <.0001 |
| C. mixo – C. het <> E. mixo – C. het | -2.6370 | 0.173 | 29.7 | -15.267 | <.0001 |
| C. mixo – E. het <> E. mixo – E. het | -1.3631 | 0.156 | 30.8 | -8.729 | <.0001 |
| C. mixo – E. het <> E. mixo – C. het | -2.6823 | 0.156 | 30.8 | -17.177 | <.0001 |
| E. mixo – E. het <> E. mixo – C. het | -1.3192 | 0.154 | 29.7 | -8.539 | <.0001 |
| **L = continuous, P = pulsed** |  |  |  |  |  |
| C. mixo – C. het <> C. mixo – E. het | -0.3546 | 0.173 | 29.7 | -2.053 | 0.1920 |
| C. mixo – C. het <> E. mixo – E. het | -1.2170 | 0.173 | 29.7 | -7.046 | <.0001 |
| C. mixo – C. het <> E. mixo – C. het | -2.2986 | 0.173 | 29.7 | -13.308 | <.0001 |
| C. mixo – E. het <> E. mixo – E. het | -0.8624 | 0.154 | 29.7 | -5.582 | <.0001 |
| C. mixo – E. het <> E. mixo – C. het | -1.9440 | 0.154 | 29.7 | -12.584 | <.0001 |
| E. mixo – E. het <> E. mixo – C. het | -1.0816 | 0.154 | 29.7 | -7.001 | <.0001 |
| **L = fluctuating, P = pulsed** |  |  |  |  |  |
| C. mixo – C. het <> C. mixo – E. het | -0.4257 | 0.156 | 30.8 | -2.726 | 0.0488 |
| C. mixo – C. het <> E. mixo – E. het | -1.5269 | 0.154 | 29.7 | -9.883 | <.0001 |
| C. mixo – C. het <> E. mixo – C. het | -1.8133 | 0.154 | 29.7 | -11.737 | <.0001 |
| C. mixo – E. het <> E. mixo – E. het | -1.1012 | 0.156 | 30.8 | -7.052 | <.0001 |
| C. mixo – E. het <> E. mixo – C. het | -1.3876 | 0.156 | 30.8 | -8.886 | <.0001 |
| E. mixo – E. het <> E. mixo – C. het | -0.2864 | 0.154 | 29.7 | -1.854 | 0.2693 |

### Experimental set-up and design

The chemostat system used consisted of 24 culture vessels (culture volume 900 ml) and corresponding medium and waste containers, tubing and peristaltic pumps (Ismatec, Wertheim, Germany). The medium inflow and the culture suspension outflow were established via a port in the cap of the culture vessel. A compressor provided the air pressure necessary to push the culture suspension through the outflow (Del Arco et al. 2020). Magnetic stirrers were used to keep the organisms in suspension. The dilution (flow through) rate was 0.1 d^-1^. Experimental communities grew in a modified WC medium (Guillard and Lorenzen 1972), which was nitrogen limited (120 µmol N L^-1^). According to previous experiments, *Cry* grows better if organic compounds are available, which can be supplied by adding soil extract. Half (60 µmol N L^-1^) of the N concentration in the modified WC medium, therefore, originated from a soil extract prepared following the instructions of Kies (1967). An additional 60 µmol N L^-1^ were added using the WC Nitrogen stock solution (NaNO_3_). Temperature was kept constant (18 °C), and illumination of the cultures vessels was from the side, with a light to dark cycle of 12:12 hours.

Due to material, space and time constraints, we divided the experimental investigation of the competition between mixotrophic and heterotrophic ciliates into two parts. The combinations *Euplotes octocarinatus* (*E. het*) – *Coleps viridis*. (*C. mixo*) and *Coleps hirtus* (*C. het*) – *Euplotes daidaleos* (*E. mixo*) were investigated in experiment 1 and the combinations *E. het* – *E. mixo* and *C. het* – *C. mixo* in experiment 2. We chose a 2 x 2 x 2 x 3 design for the experiments, exposing two different species combinations to two light regimes (constant and fluctuating) and two regimes of prey supply (continuous and pulsed), each of those replicated three times, resulting in 24 experimental units per experiment. The initial total ciliate biovolume was 10.76 x 10^5^ µm^3^ ml^-1^ in experiment 1 and 6.30 x 10^5^ µm^3^ ml^-1^ in experiment 2, with each of the ciliate species being initially incubated at equal biovolume. Constant light was supplied at an intermediate intensity (33 µmol m^-2^ s^-1^ photosynthetic photon flux density (PPFD)). Under fluctuating light, phases of high (59 µmol m^-2^ s^-1^ PPFD) and low light (7 µmol m^-2^ s^-1^ PPFD) alternated at the interval length of 8 days, resulting in an equal amount of light provided over the course of the experiments under both constant and fluctuating light conditions. To establish the experimental light intensities, the PPFD was determined inside of the empty culture vessels. Intermediate and low light intensities were realized by covering the culture vessels of the chemostat systems by black netting of different mesh size. The microalgal prey was added manually. To ensure the cells were in a comparable condition each time, we used dense batch cultures of *Cry* that were in or close to steady state condition. Abundance was determined microscopically (see below), before adding the appropriate volume of culture suspension to the chemostats. Continuous prey supply consisted of 1000 cells/ml of *Cry* every day, while under pulsed prey supply 8000 cells/ml were fed every eighth day. This way the time average of light intensity and prey supply was the same in all treatments. Duration of the experiments was 48 days.

Experimental units were sampled every second day using a hypodermic syringe and cannula (1.0 x 200 mm, BD Plastipak, B. Braun, Melsungen, Germany; neoLab Migge, Heidelberg, Germany). The total sample volume (60 ml) was subsampled as follows: Subsamples for microscopic analyses (30 ml) of ciliate and microalgal abundance were taken every second day, and subsamples for Glutaraldehyde fixation (9 ml) every fourth day.

### Sample processing and analysis

Samples for microscopic analysis were fixed with Lugol’s solution (1% final concentration) and stored in brown glass bottles. *Cry* concentration was analysed microscopically (Leica DMIL) using a subsample of 1 ml. If possible, at least 400 cells per sample were counted in grids at 100x magnification (Lund et al. 1958). In cases where algal concentrations were too low following this method, two 0.5 mm transects at 100x magnification (equalling a sixth of the counting chamber or a subsample of 0.335 ml) were counted. Ciliate concentrations were counted in a subsample sized 2 or 3 ml. If no ciliates were detected, a concentration of 0.5 x the detection limit (0.25 cell ml^-1^) was assumed (Clarke 1998). Since *C. het* and *C. mixo* were indistinguishable by light microscopy, total *Coleps* concentrations were quantified in the *C. mixo* – *C. het* treatment. We determined the *C. mixo* to *C. het* ratio via epifluorescence microscopy (Axiophot, Zeiss) using Glutaraldehyde preserved samples (final concentration 1 %). Here, *C. mixo* could be distinguished from *C. hetero* under blue light excitation by the presence of algal symbionts, which could be clearly distinguished from potentially ingested algae of *C. hetero* at 1000x magnification. The *C. mixo* and *C. hetero* proportions determined were then used to calculate the concentrations of the different *Coleps* species. The dimensions of 20 individuals of each ciliate and algal species were initially determined using a digital image system program (Cell-P) to calculate the average specific biovolume (Hillebrand et al. 1999). These data were used to calculate population biovolume throughout the experiments.

### Data Analyses

In experiment 1, contaminations with *E. mixo* occurred in one of the replicates of the *E. het* – *C. mixo* combination that received fluctuating light and continuous prey supply, and in one replicate of the *E. het* – *C. mixo* combination with fluctuating light and pulsed prey supply. Therefore, results from these replicates covering the last full light cycle (days 33 – 48) had to be removed from the data set. In experiment 2, two replicates were lost from the species combination *C. het* – *C. mixo*, i.e. one in the treatment that had received constant light and pulsed prey and the other one in the treatment fluctuating light and continuous prey supply.

### Community Dynamics

We used a linear mixed model ANOVA to analyse community dynamics of both experiments, where the log-biomass ratio of mixotrophic and heterotrophic ciliates served as response variable. Species combination, and modes of light and prey supply were entered as factors, time (days) was set as trend and as factor for random fluctuations over time. Model assumptions were validated graphically. The analysis (Type III ANOVA) was performed with R version 3.6.3 (R Development Core Team 2020) using RStudio version 1.2.5042 (RStudio, Boston, USA), using R-package lme4 (Bates et al. 2015) with Satterthwaite’s method of degrees of freedom in combination with lmerTest (Kuznetsova et al. 2017). Post-hoc tests (contrasts) used Kenward-Roger method for degrees-of-freedom method and Tukey P-value adjustment method for comparing a family of 4 estimates.

### Species similarity and rate of competitive exclusion

To estimate the rate of competitive exclusion of the inferior competitor in our ciliate communities, we fitted a linear regression model to the time series of the logarithm of the relative proportion of the inferior competitor to the total ciliate biovolume. The slope of the linear regression model was used as an estimate of the extinction rate of the inferior competitor. To determine whether this extinction rate was potentially related to the similarity of the two different competitors, we additionally assessed the similarity or dissimilarity between a pair of species based on a synchronous or asynchronous course in the short-term fluctuations of their population dynamics as well as the difference in the temporal variation or amplitude of their short-term fluctuations. The synchronicity in the short-term fluctuations of the species population dynamics was estimated based on the pairwise *Pearson* correlation coefficient *r* between the detrended logarithmic biovolumes of the different species. A detrending of the logarithmic biovolumes was achieved by fitting a linear regression model to the time series of the logarithmic biovolumes and subsequently subtracting the resulting biovolume estimates of the linear model from the original measurements at the corresponding time points. While positive values of r indicate that both competitors are responding similar to the current environmental conditions, negative values of r indicate the opposite. Hence, the two species were assumed to be ecologically more similar the more positive the corresponding correlation coefficient r was between their short-term fluctuations in the population dynamics. In addition, we evaluated whether two competitors differed in the amplitudes of the short-term fluctuations of their population dynamics based on the log-response ratio between the standard deviations of the species’ detrended logarithmic biovolumes. A log-response ratio of 0 indicates that there is no difference between the species in the short-term variation of their biovolumes. In contrast, the larger the absolute value of the log-response ratio, the more different the species are in respect to the short-term variation of their logarithmic biovolumes. Overall, we assumed two species to be more similar the more synchronous they responded in their short-term population dynamics and the lower the difference was in the variation of their short-term fluctuations. The corresponding data analyses and plotting were performed using *MATLAB*, version 9.10 (MATLAB, 165 2019. version 9.6.0 (R2019a), Natick, Massachusetts: The MathWorks Inc, 2021).

## RESULTS

### Trajectories of heterotrophic and mixotrophic ciliates and their microalgal prey

Although population dynamics were affected by both light fluctuations and prey pulses, in each species combination one species gained dominance at the end of the experiment (Fig. 1). In the same genus combination *C. het – C. mixo* it was the heterotroph, while the mixotroph became dominant in the *E. het* – *E. mixo* combination irrespective of treatment combination. In the genus mixtures, the heterotroph dominated the combination *E. het* – *C. mixo* at the end of the experiment, whereas it was the mixotroph in the *C. het* – *E. mixo* combination.

**Figure 1:**
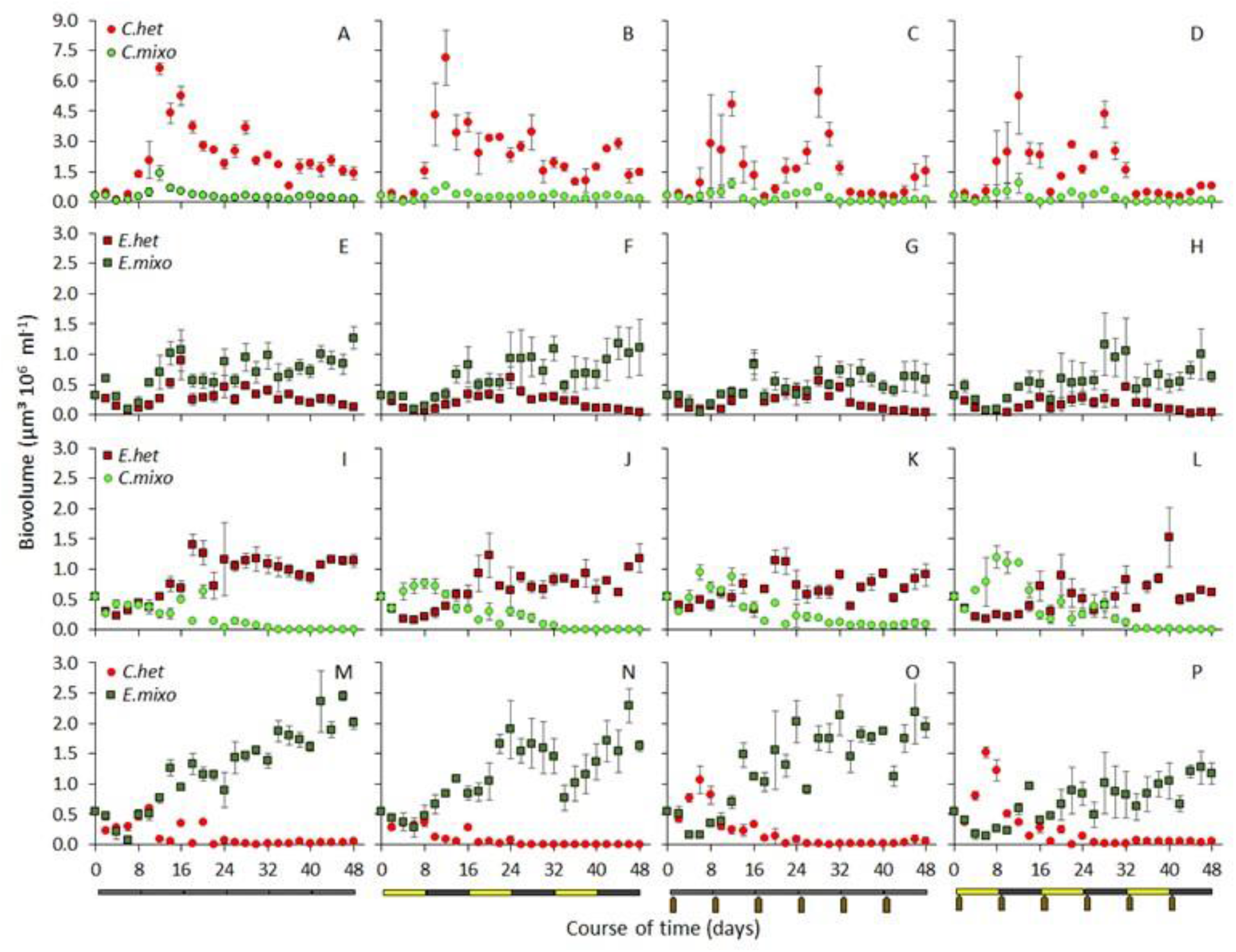
Population dynamics of heterotrophic and mixotrophic ciliates over the course of time, shown as biovolume development. Species combinations: *C. het* – *C. mixo* (A-D), *E. het* – *E. mixo* (E-H), *E. het* – *C. mixo* (I-L), *C. het* – *E. mixo* (M-P). Resource supply modes: continuous light and constant prey (A, E, I, M), fluctuating light and constant prey (B, F, J, N), continuous light and pulsed prey (C, G, K, O), fluctuating light and pulsed prey (D, H, L, P).

#### Same genus combinations

##### C. het – C. mixo

The temporal pattern of *C. mixo* biovolume basically followed the one of *C. het* on a lower level regardless of resource supply mode. Biovolume increased quickly, showing peaks on day 12 and 28 (Fig. 1A-D). At constant prey supply, a pronounced first *C. het* biovolume peak was followed by a smaller one (Fig. 1A, 1B). A third peak was observed on day 44, when constant prey supply was combined with fluctuating light (Fig. 1B). While *C. het* biovolume remained on a high level at constant prey supply, *C. mixo* biovolume remained low from day 18 onwards (Fig. 1A, 1B). The corresponding biovolume of the microalgal prey *Cry* in these treatments increased until day 6, then declined and fluctuated slightly (Fig 2A, 2B), indicating that ciliate density was high enough to keep algal prey densities at low levels close to depletion. Under pulsed prey supply, *C. het* biovolume peaks on day 12 and 28 were in the same concentration range, while in-between the peaks the biovolume was much lower compared to constant resource supply and remained low until the end of the experiment (Fig. 1C, 1D). Biovolume peaks of *C. mixo* were notably lower (Fig. 1C, 1D). *Cry* biovolume decreased after the pulses; the decreases were especially pronounced before prey pulses 3, 5 and 6, indicating phases of food shortage (Fig. 2C, 2D).

**Figure 2:**
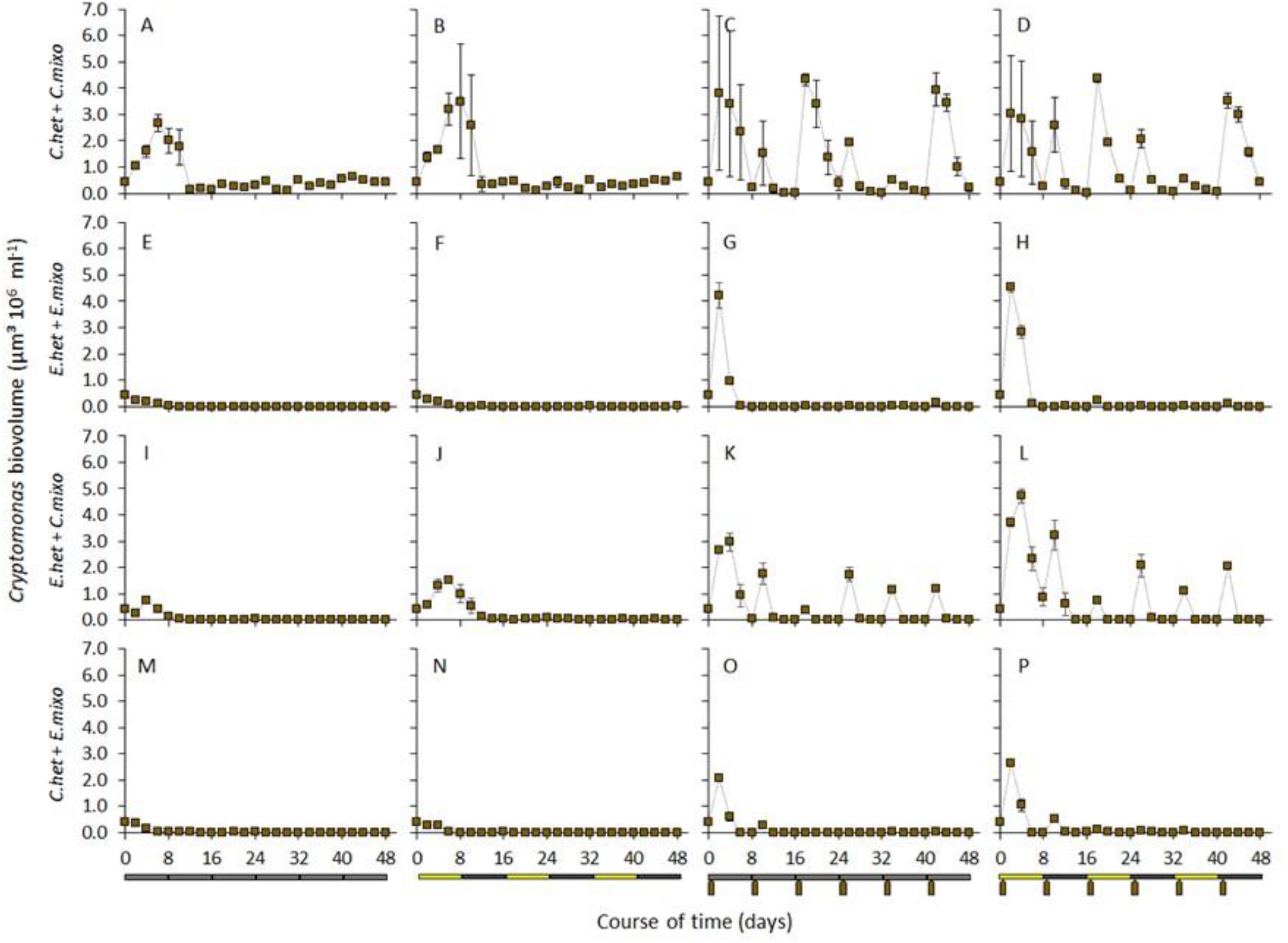
Time course of microalgal prey biovolume. Species combinations: *C. het* – *C. mixo* (A-D), *E. het* – *E. mixo* (E-H), *E. het* – *C. mixo* (I-L), *C. het* – *E. mixo* (M-P). Resource supply modes: continuous light and constant prey (A, E, I, M), fluctuating light and constant prey (B, F, J, N), continuous light and pulsed prey (C, G, K, O), fluctuating light and pulsed prey (D, H, L, P).

##### E. het – E. mixo

During the first week of the experiment, biovolume of *E. het* and *E. mixo* declined in all treatments (Fig 1E-H). After this initial dip, both populations grew with *E. mixo* displaying a slightly faster increase in biovolume than *E. het*. Biovolume of *E. mixo* peaked (Fig. 1E-G) or was increased (Fig. 1H) on day 16. Under continuous light (Fig. 1E, 1G) *E. het* biovolume peaked also on day 16, while a later peak (day 24) was observed when light fluctuated and prey supply was constant (Fig. 1F). From day 16 on, biovolume of both *E. het* and *E. mixo* fluctuated in all treatments until around day 32, where *E. het* biovolume started to decline while the one of *E. mixo* increased or remained on the same level (Fig. 1E-H). Under continuous light and pulsed prey supply, *E. het* and *E. mixo* biovolume was comparable from the start of the experiment to day 32 (Fig. 1G). In the other resource supply treatments *E. mixo* biovolume remained above the one of *E. het* throughout the experiment (Fig. 1E, 1F, 1H). Under constant prey supply, *Cry* biovolume decreased during the first days of the experiment and remained around detection level from day 8 onwards (Fig. 2E, 2F), indicating strong grazing pressure. Pulsed prey supply led to an initial increase in *Cry* biovolume (Fig. 2G, 2H), which was followed by a decrease reaching detection level on day 8. Later in the experiment *Cry* biovolume approached detection level two days after a prey pulse was given, indicating severe resource shortage.

#### Mixed-genus combinations

##### E. het – C. mixo

*E. het* biovolume decreased during the first days of the experiment, while *C. mixo* biovolume started increasing from day 2. This initial dip in *E. het* biovolume was much longer and more pronounced under fluctuating light conditions (Fig. 1 J, L), where *C.mixo* gained a clear dominance in the first 14 days of the experiment as opposed to constant light (Fig. 1 I, K). This pattern coincided with increases in *Cry* biovolume during the first high light cycle (Fig. 2J, 2L) and was more pronounced with pulsed prey supply (Fig. 1L). Under continuous light and constant prey supply *C. mixo* biovolume was initially only marginally higher than that of *E. het* (Fig. 1I), and this difference was a bit more pronounced for the first 12 days when continuous light was combined with pulsed prey supply (Fig. 1K). After the initial increase, *C. mixo* biovolume showed some fluctuations under all resource supply modes and then declined. Except for the continuous light and pulsed prey supply treatment (Fig. 1K), *C. mixo* biovolume was close to the detection limit during the last 14 day of the experiment. After the initial dip, *E. het* biovolume increased and remained on a high level, which was most pronounced at continuous light and prey supply. Pulsed prey supply was connected to fluctuations in *E. het* biovolume (Fig. 1O, 1P). After an initial increase, *Cry* biovolume (Fig. 2I, 2J) stayed at a low level under constant prey supply, indicating that grazing pressure increased quickly and remained high until the end of the experiment. Under pulsed prey supply (Fig. 2K, 2L), *Cry* biovolume was depleted within four days from the second pulse onwards, indicating phases of resource shortage.

##### C. het – E. mixo

*E. mixo* biovolume declined during the first days of the experiment and during this time *C. het* biovolume increased (Fig. 1M-P). Peaks in *C. het* biovolume (day 6) were observed under fluctuating prey supply, with the peak being more pronounced when pulsed prey supply was combined with fluctuating light (Fig. 1P). After an initial phase of similarity (Fig. 1N) or *C. het* dominance (Fig. 1O, 1P), *C. het* declined while *E. mixo* biovolume increased and became dominant regardless of resource supply mode. This dominance was least pronounced with pulsed prey supply and fluctuating light (Fig. 1P). Under constant prey supply *Cry* biovolume (Fig. 2M, 2N) decreased and remained on a low level from day 6, indicating high grazing pressure. Except for the first pulse, where depletion occurred on day 6, *Cry* biovolume (Fig. 2P) decreased rapidly after prey pulses, indicating phases of severe resource shortage.

### Proportion of mixotrophic and heterotrophic ciliates

As illustrated by the log-ratio (Fig. 3), the proportion of mixotrophic and heterotrophic ciliates was more strongly affected by species combination than by resource supply treatments. In all combinations of light and prey supply modes, mixotrophic ciliates gained dominance by the end of the experiment when *E. mixo* was part of the species combination, while heterotrophic ciliates gained dominance when *C. mixo* was present. That means that in all two-genus combinations, *Euplotes* was superior over *Coleps* at the end of the experiment irrespective of nutritional strategy (heterotrophic versus mixotrophic). Both species combination and prey supply significantly altered biomass log-ratios of mixotrophs/heterotrophs (significant main effects, mixed model ANOVA, Table 1), while light supply effects depended on consumer combination (significant 2-way interaction, Table 1) and also varied with prey supply and time (significant three-way interactions, Table 1). Accordingly, ciliate dominance patterns changed over time in dependence of light and prey supply (Fig. 3). The degree of dominance, however, differed among species combinations and was stronger in two-genus combinations (*Euplotes* combined with *Coleps*) than in same genus combinations (*E. mixo* – *E. het*, *C. mixo* – *C. het*). These differences were significant regarding time averaged log ratios, except for the species combinations *C. het* – and *E. het* – *C. mixo* in all but the fluctuating light – pulsed prey treatment, and the combinations *E. het* – *E. mixo* and *C. het* – *E- mixo* under fluctuating light and pulsed prey supply (significant contrasts, Table 1).

**Figure 3:**
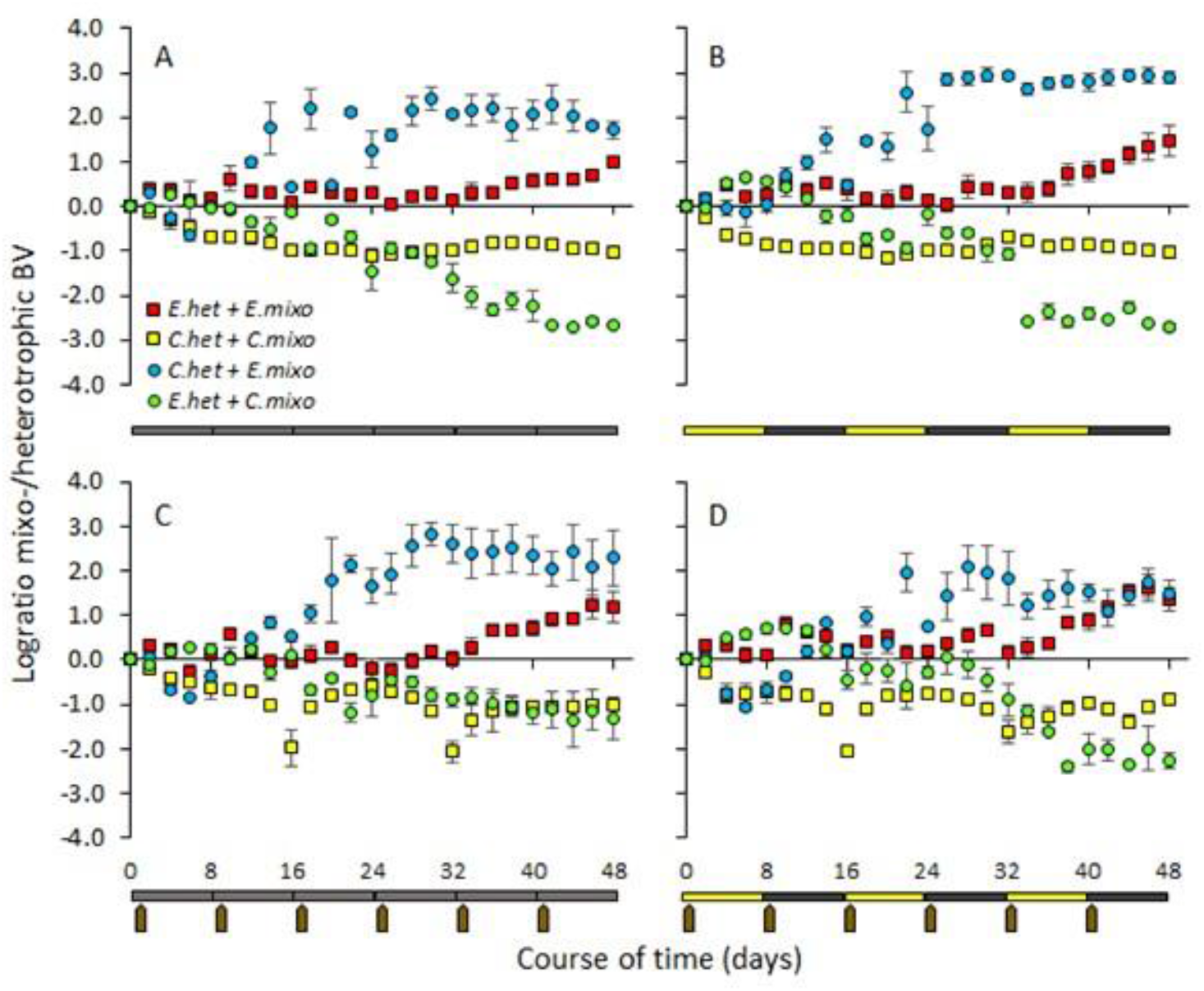
Proportion of mixotrophic to heterotrophic biovolume (BV) over the course of time, expressed as log-ratio (log10(mixotrophic BV/heterotrophic BV). Resource supply modes: continuous light and constant prey (A), fluctuating light and constant prey (B), continuous light and pulsed prey (C), fluctuating light and pulsed prey (D).

The dominance of *E. mixo* at the end of the experiment was more pronounced in combination with *C. het* than in combination with *E. het* (Fig. 3). Externally forced cycles were especially observed in the dominance patterns of the *E. het* – *C. mixo* combination under fluctuating light (Fig. 3B, D). *C. mixo* gained dominance during the first high light phase, decreased in biomass proportion during the dark phase, and increased again in the following high light phase, even though it was not dominant anymore. At the end of the experiment, however, *E. het* dominated all resource supply treatments in combination with *C. mixo*, which was least pronounced under constant light and fluctuating prey (Fig. 3C). In the *C. het* – *C. mixo* combinations, *C. het* periodically dominated strongly under pulsed prey supply, whereas dominance patterns remained un-changed under constant prey supply. In the *E. het* – *E. mixo* combinations, biovolume proportions were similar and fluctuated slightly, especially in response to prey pulses, before *E. mixo* clearly gained dominance towards the end of the experiment. These complex patterns are well reflected in the significant four-way interaction between species combination, light and prey supply modes, and time (mixed model ANOVA, Table 1).

### Species similarity and rate of competitive exclusion

Overall, species identity affected the outcome of ciliate competition much more than the applied resource supply regimes. While competitive exclusion occurred rather slowly in our treatments that comprised either the two different *Coleps* strains or the two different *Euplotes* strains, the estimated extinction rate of the inferior competitor was relatively high (indicated by more negative values) in the treatments that comprised a combination of *Coleps* and *Euplotes* (Fig. 4A). In addition, the short-term fluctuations in the population dynamics of the ciliates were strongly positively correlated and showed lower differences in their temporal variation in the treatments that comprised the two different *Coleps* strains or the two different *Euplotes* strains (Fig. 4B, C). In contrast, mixed genus combinations of *Coleps* and *Euplotes* exhibited uncorrelated or negative short-term fluctuations in population dynamics with larger differences in their temporal variation. Consequently, a higher extinction rate of the inferior competitor was associated with more asynchronous short-term fluctuations in the population dynamics (*Pearson correlation*: r = 0.58, p < 0.0001, Figure 4D) and larger differences in the short-term temporal variation of the population dynamics (*Pearson correlation*: r = −0.79, p < 0.0001, Figure 4E). Our results therefore suggest that competitive exclusion occurred the faster the more differently the species responded to the environmental conditions and thus the more dissimilar the species were.

**Figure 4:**
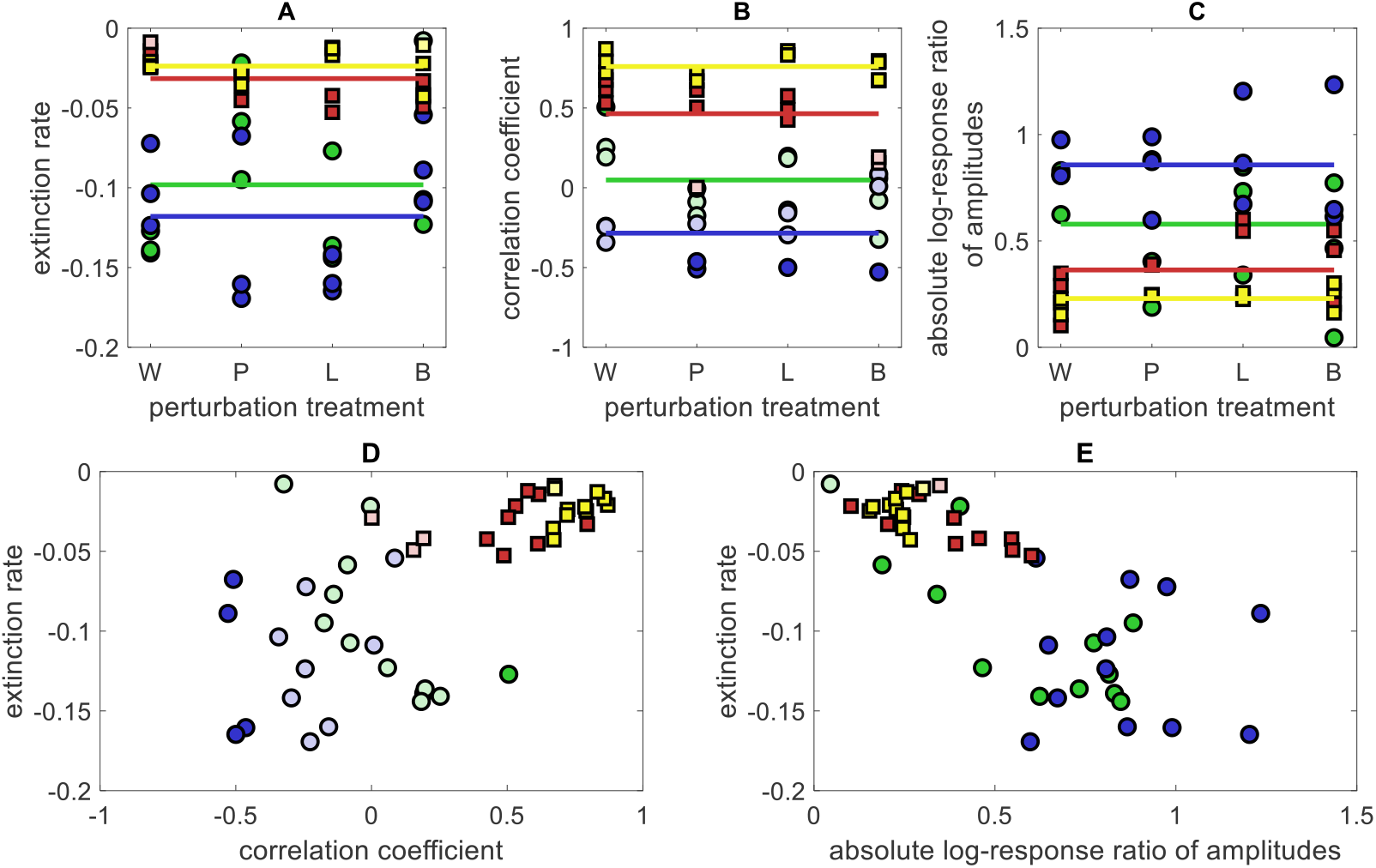
Relationship between extinction risk and trait similarity. The upper panels show the extinction risk of the inferior competitor (A), the temporal correlation between the short-term fluctuations of the two competitors (B) and the absolute value of the log-response ratio of the amplitudes, i.e. standard deviation, of the short-term fluctuations of the competitors (C) in dependence of the different resource supply modes, i.e. continuous light and constant prey (W); continuous light and pulsed prey (P), fluctuating light and constant prey (L), fluctuating light and pulsed prey (B). The horizontal lines indicate the average effect sizes for the different species combinations across the different resource supply modes. The lower panels show the relationship between the estimated extinction rate of the inferior competitor and the estimated correlation coefficient between the competitors’ short-term fluctuations (D) respectively the difference between the estimated amplitudes of the competitors’ short-term fluctuations (E). The colors indicate the different species compositions, i.e. *C. het* – *C. mixo* (yellow), *E. het* – *E. mixo* (red), *E. het* – *C. mixo* (green), *C. het* – *E. mixo* (blue). Regarding panel (B) and (D), dark-colored dots indicate significant (p<0.05) correlation coefficients whereas light-colored dots represent insignificant (p>0.05) correlation coefficients.

## DISCUSSION

Competition between mixotrophic and heterotrophic ciliates in our experiment was strongly affected by species composition. Irrespective of light and prey supply, *Euplotes* dominated both mixed genus combinations by the end of the experiment, i.e. *E. het* in the *E. het* – *C. mixo* combination and *E. mixo* in the *C. het* – *E. mixo* combination. In the same genus combinations, the heterotroph gained dominance in the *C. het* – *C. mixo* combination, whereas it was the mixotroph in the *E. het* – *E. mixo* combination. Overall, genus-specific differences in response to resource supply mode led to faster competitive exclusion in mixed-genus combinations than in more similar species of the same genus. While fluctuations in resource supply did not alter the qualitative outcome of species competition, it influenced the shape of the population dynamics throughout the experiment and promoted species coexistence in some species combinations where the inferior competitor was reaching higher biomasses at the end of the experiment.

### Effects of resource supply mode

While the resource supply mode influenced ciliate population dynamics, it did not alter the qualitative outcome of species competition in the different species combinations. Regardless of the light regime, the heterotroph dominated the same genus combination *C. het* – *C. mixo* at constant prey supply. Following the trajectory of the heterotroph on a lower biovolume level, the mixotroph persisted at low proportions. In contrast to our expectations, prey pulses entailing periods of resource depletion did not promote the mixotroph, but induced a stronger growth response and dominance in the heterotroph reflected in cyclic biovolume shifts in response to prey pulses irrespective of light regime. This is in line with a preliminary experiment, where *C. het* exhibited a higher maximum specific ingestion rate for the microalgal prey compared to *C. mixo* (unpublished data), indicating that *C. het* was able to use a larger proportion of the prey supplied for population growth. Similarly, *Coleps hirtus viridis* without algal symbionts displayed much higher ingestion and feeding rates than the same strain with algal symbionts (Auer et al. 2004). While the proportion of prey acquired by competing species may affect their dominance in a community, the competitor with the lowest resource threshold level should be superior and exclude other species over time (Rothhaupt 1988, Weisse 2017). Light availability was expected to alter the prey threshold required by *C. mixo* based on its ability to supplement its energy supply with photosynthetic carbon fixation. At constant light conditions or in high light phases, a lower prey threshold level in *C. mixo* should have therefore promoted *C. mixo* growth. However, *C. mixo* biovolume levels remained low irrespective of light regime and *C. het* remained the dominant competitor. This indicates that the selected light conditions in our experiment might not have been sufficient to balance the costs of hosting symbionts via photosynthesis and to sufficiently complement phagotrophic nutrition with photosynthetic production allowing enhanced growth. Whether light fluctuations are generally beneficial or detrimental for mixotrophic growth will further depend on the curvature of the mixotrophs’ light-dependent growth function and the simultaneous effects on the growth of the phototrophic prey that serves not only as an alternative food source, but also as a competitor for shared nutrients and light. In general, ciliates have the potential to reach higher gross growth efficiencies through acquired phototrophy (Stoecker et al. 2009, Leles et al. 2018), in particular when combined with sufficient food supply. For example, an increased food uptake of the eSNCM *P. bursaria* during phases of high light intensity has been interpreted as a strategy to increase nutrient supply to the symbionts to meet their demands during phases of increased photosynthetic activity (Berk et al. 1991). Similarly, we observed an increase in the relative abundance of *C. mixo* when prey pulses were associated with either constant light or high light phases, especially when heterotroph abundances were low, while being unaffected or even declining when prey pulses coincided with low-light conditions. As prey pulses led to stronger oscillations in the *C. het* population than in *C. mixo*, we assume that the resource-dependent growth function of *C. het* may be much steeper than the one of *C. mixo* resulting in more pronounced consumer-prey cycles for *C. het*. Overall, despite being inferior to its heterotrophic counterpart, the mixotroph was able to withstand the resource shortage caused by *C. het*., and coexisted at a low level with the heterotroph through niche expansion via its acquired metabolism through symbionts (Hsu and Moeller 2021, Hsu et al. 2022).

In the other same-genus combination, *E. mixo* and *E. het*, both species which have been shown to express similar specific ingestion rates (preliminary experiment, unpublished data) coexisted at similar levels for the first 32 days of the experiment. However, towards the end of the experiment, the mixotroph gained dominance, which was a bit more pronounced under fluctuating light supply compared to constant light. Apparently, the high light phases in combination with low light phases were more beneficial for *E. mixo* compared to constant intermediate light conditions for the increased nutritional benefit through acquired phototrophy to play out. Preliminary observations showed that *E. mixo* is also able to grow in the dark when sufficient prey is provided (unpublished data). The capacity to potentially grow heterotrophically in the dark and to complement heterotrophy with photosynthetic gains during the day likely led to competitive dominance of *E. mixo.* This was different in an obligate ciliate GNCM which exhibited similar overall growth rates compared to its heterotrophic counterpart despite the energy subsidy received via photosynthesis, as mixotrophic growth was not possible in the dark (Schoener and McManus 2017). Lowe et al. (2016) indicated for the eSNCM *P. bursaria* that its symbiosis changed from costly to beneficial with increasing light intensity, leading to higher growth rates compared to symbiont-free *P. bursaria*. This supports the idea that mixotrophic performance is likely determined by specific irradiance levels. In our experiment, high and intermediate light levels were apparently sufficient to promote *E. mixo* over its heterotrophic competitor despite similar grazing efficiencies, while for *C. mixo* this was not the case. Higher ingestion rates therefore allowed *C. het* to gain and maintain dominance.

In both mixed-genus combinations (*E. het* – *C. mixo, C. het* – *E. mixo), Euplotes* gained dominance likely because of their higher ingestion rates of microalgal prey and their ability to efficiently utilize bacteria as an additional prey in our non-axenic cultures, which has been observed in preliminary culture experiments (i.e. both *Euplotes* strains can grow on a bacterial diet, while the *Coleps* strains cannot, unpublished data). In the *C. het* – *E. mixo* combination, prey pulses entailing periods of resource depletion did not promote *E. mixo* at first as expected, but promoted the growth of *C. het* instead, in particular at the beginning of the experiment. This may hint towards a stronger curvature in the resource-dependent growth function of *E. mixo* making it more sensitive to resource fluctuations than *C. het.* However, *E. mixo* gained dominance, nevertheless, and produced higher biovolume levels in the *C. het* – *E. mixo* combination than *E. het* in the *E. het* – *C. mixo* combination. This can be attributed to the higher growth efficiency of *E. mixo* that it acquired via phototrophy (Stoecker et al. 2009, Leles et al. 2018). In both combinations, *Coleps* exhibited an immediate growth response, while *Euplotes* went through a pronounced lag-phase at the beginning of the experiment, enabling *Coleps* to effectively use the inactivity of its competitor for grazing and population growth (Flöder et al. 2021). When starting to grow, however, the high grazing efficiency of *Euplotes* created strong prey shortages for *Coleps*, promoting its competitive exclusion. However, persistence of *C. mixo* at the end of the experiment was promoted by pulses in the prey supply, which might be attributed to a higher starvation resistance of *C. mixo* at periods of food shortage. In contrast, persistence of *C. hetero* also required pulses in the light supply which might have generated windows of opportunities for *C. hetero* in which it has a competitive advantage.

Overall, in contrast to our expectations, coexistence of heterotrophs and mixotrophs were not generally promoted by fluctuating / pulsed resource supply; only *E. mixo* and *C. het* exhibited the greatest coexistence at both fluctuating light and pulsed prey, while the other combinations showed highest coexistence at different specific environmental conditions.

### Effects of species composition – functional traits and extinction rates

While *Coleps* and *Euplotes* have a similar prey spectrum, both feeding on planktonic algae and bacteria (Fenchel 1986, Madoni et al. 1990, Früh et al. 2011, Lawrence and Snyder 2011, Buonanno et al. 2014), other functional traits of the organisms differ such as their maximum specific ingestion rate, the ability of *Euplotes* to exclusively grow on bacteria, and the specific light conditions promoting mixotrophic nutritional gains (see above). These differences may explain the general dominance patterns observed in our competition experiment. In addition, other traits might have influenced population dynamics over the course of the experiment. For instance, potentially higher metabolic rates of the smaller *Coleps* compared to the larger *Euplotes* (Savage et al. 2004, Moorthi et al. 2016) could have led to a quicker response of both, *C. het* and *C. mixo* to a high supply of microalgal prey in the pulsed treatments. In this context, the observed longer lag-phase of *E. het* and *E. mixo* at the beginning of the experiment likely also played a role, offering a window of opportunity for fast growing *Coleps* strains. This extended lag phase has also been reported for other *Euplotes* species (Garstecki and Wickham 2001, Ning et al. 2011). *Cry* biovolume in mixed-genus combinations was reduced to similarly low levels as in the *E. het* – *E. mixo* combination, while the remaining prey biovolume in the *C. het* – *C. mixo* combination was higher. This indicates that *E. het* and *E. mixo* have lower resource threshold levels and, therefore, are better competitors for *Cry* than *C. het* and *C. mixo*. This observation is confirmed by results of Cadotte et al. (2006) who ranked the competitive ability of *Euplotes* higher than the one of *Coleps*. The results of our experiment indicate that functional traits determining ciliate feeding and growth responses were more similar in organisms of the same genus than for organisms of the functional groups ‘heterotrophs’ and ‘mixotrophs’, supporting our initial expectation regarding the effects of resource supply regime on same-genus versus mixed-genus combinations. According to classical theory, the likelihood of species coexistence strongly depends on the similarity of competitors with respect to their functional traits, e.g. resource requirements, starvation and grazing resistance, and natural mortality (MacArthur & Levins 1967). Following modern coexistence theory, there are two different possibilities for species to achieve stable coexistence: either being very similar (promoting small fitness differences) or being sufficiently different (promoting large niche differences) (Chesson 2000). Competitive exclusion is expected when species exhibit trait differences that do not give rise to sufficiently large niche differences but already generate substantial fitness differences (Chesson 2000). This theoretical expectation is supported by our experimental findings where competitive exclusion occurred much faster in the treatments that comprised ciliates of different genera than in the treatments that comprised ciliates of the same genera. The degree of similarity among competing ciliates is not only supported by their phylogenetic relationship but also by the (dis)similarity in their dynamical responses measured in terms of synchronization and differences in the amplitude of their short-term fluctuations in our experiments. Hence, the outcome of competition between different ciliates in our experiment was not only driven by differences in light-harvesting and resource acquisition (feeding) traits that are associated with the presence or absence of algal symbionts, but to a larger extent by other functional differences described above. This highlights that patterns of species coexistence are rarely explained by single traits and, thus, require a multi-dimensional trait perspective. Accordingly, coexistence of mixotrophic, phototrophic and heterotrophic species cannot only be understood through a simple energetic trade-off, but requires to take into account species differences in minimum resource requirements, differences in the curvatures of species’ resource-dependent growth functions and, thus, sensitivities to resource fluctuations as well as direct and indirect species effects on the species’ fitness. For example, a heterotrophic species may not only negatively affect its mixotrophic competitor by feeding on its shared phototrophic prey but may also positively affect mixotrophic growth by reducing the negative effects of the phototrophic prey caused by competition for shared nutrients and light.

Our results therefore emphasize that genus- / species-specific traits other than related to nutritional mode may override the relevance of acquired phototrophy by heterotrophs in competitive interactions. The advantage of photosynthetic carbon fixation of symbiont-bearing mixotrophs in competition with pure heterotrophs may differ greatly among different mixotrophs, playing out under different environmental conditions and depending on the specific requirements of the species.

## ACKNOWLEDGEMENTS

We would like to thank Christian Spindler and Ronny Steinberg for their help with the experimental work. This study was funded by the German Research Foundation (DFG, DynaTrait: MO 1931/4-1)

## AUTHOR CONTRIBUTIONS

Stefanie Moorthi, Toni Klauschies and Sabine Flöder conceived the study and conjointly wrote the manuscript. Sabine Flöder, Moritz Klaassen, Tjardo Stoffers and Max Lambrecht conducted chemostat experiments, sample processing and analyses. Sabine Flöder and Toni Klauschies performed the data analyses. All authors approved of the submission of the manuscript.

## Open Research statement

The data that support the findings of this study are available on request from the corresponding author, S.F. However, since the data enclosed in the manuscript have not yet been published elsewhere, all authors have agreed on making them only fully available after acceptance. We will use Dryad to permanently archive experimental data and R-code. The R-code used is not novel. All code is properly cited and publicly available.

## REFERENCES

Andersen, K. H., D. L. Aksnes, T. Berge, Ø. Fiksen, and A. Visser. 2015. Modelling emergent trophic strategies in plankton. Journal of Plankton Research 37:862–868.

Archibald, J. M. 2009. The puzzle of plastid evolution. Current Biology 19:R81–R88.

Auer, B., E. Czioska, and H. Arndt. 2004. The pelagic community of a gravel pit lake: Significance of Coleps hirtus viridis (Prostomatida) and its role as a scavenger. Limnologica 34:187–198.

Bates, D., M. Mächler, B. Bolker, and S. Walker. 2015. Fitting Linear Mixed-Effects Models Using lme4. Journal of Statistical Software 67:1–48.

Berk, S. G., L. H. Parks, and R. S. Ting. 1991. Photoadaptation alters the ingestion rate of Paramecium bursaria, a mixotrophic ciliate. Applied and environmental microbiology 57:2312–2316.

Buonanno, F., A. Anesi, G. Guella, S. Kumar, D. Bharti, A. La Terza, L. Quassinti, M. Bramucci, and C. Ortenzi. 2014. Chemical offense by means of toxicysts in the freshwater ciliate, Coleps hirtus. Journal of Eukaryotic Microbiology 61:293–304.

Burkholder, J. A., P. M. Glibert, and H. M. Skelton. 2008. Mixotrophy, a major mode of nutrition for harmful algal species in eutrophic waters. Harmful Algae 8:77–93.

Cadotte, M. W., D. V. Mai, S. Jantz, M. D. Collins, M. Keele, and J. A. Drake. 2006. On testing the competition-colonization trade-off in a multispecies assemblage. The American Naturalist 168:704–709.

Calbet, A., M. Bertos, C. Fuentes-Grünewald, E. Alacid, R. Figueroa, B. Renom, and E. Garcés. 2011. Intraspecific variability in Karlodinium veneficum: growth rates, mixotrophy, and lipid composition. Harmful Algae 10:654–667.

Caron, D. A., P. D. Countway, A. C. Jones, D. Y. Kim, and A. Schnetzer. 2012. Marine protistan diversity. Annual review of marine science 4:467–493.

Caron, D. A., R. W. Sanders, E. L. Lim, C. Marrasé, L. A. Amaral, S. Whitney, R. B. Aoki, and K. Porter. 1993. Light-dependent phagotrophy in the freshwater mixotrophic chrysophyte Dinobryon cylindricum. Microbial Ecology 25:93–111.

Clarke, J. U. 1998. Evaluation of Censored Data Methods To Allow Statistical Comparisons among Very Small Samples with Below Detection Limit Observations. Environmental Science & Technology 32:177–183.

Del Arco, A., N. Woltermann, and L. Becks. 2020. Building up chemostats for experimental ecoevolutionary studies. protocols.io.

Dolan, J. R., and M. T. Pérez. 2000. Costs, benefits and characteristics of mixotrophy in marine oligotrichs. Freshwater Biology 45:227–238.

Edwards, K. F. 2019. Mixotrophy in nanoflagellates across environmental gradients in the ocean. Proceedings of the National Academy of Sciences 116:6211–6220.

Esteban, G. F., T. Fenchel, and B. J. Finlay. 2010. Mixotrophy in Ciliates. Protist 161:621–641.

Fenchel, T. 1986. Protozoan filter feeding. Prog. Protistol. 1:65–113.

Figueroa-Martinez, F., A. M. Nedelcu, D. R. Smith, and A. Reyes-Prieto. 2015. When the lights go out: the evolutionary fate of free-living colorless green algae. New Phytologist 206:972–982.

Flöder, S., L. Bromann, and S. Moorthi. 2018. Inter-and intraspecific consumer trait variations determine consumer diversity effects in multispecies predator-prey systems. Aquatic Microbial Ecology 81:243–256.

Flöder, S., J. Yong, T. Klauschies, U. Gaedke, T. Poprick, T. Brinkhoff, and S. Moorthi. 2021. Intraspecific trait variation alters the outcome of competition in freshwater ciliates. Ecology and evolution 11:10225–10243.

Flynn, K. J., A. Mitra, K. Anestis, A. A. Anschütz, A. Calbet, G. D. Ferreira, N. Gypens, P. J. Hansen, U. John, and J. L. Martin. 2019. Mixotrophic protists and a new paradigm for marine ecology: where does plankton research go now? Journal of Plankton Research 41:375–391.

Flynn, K. J., D. K. Stoecker, A. Mitra, J. A. Raven, P. M. Glibert, P. J. Hansen, E. Granéli, and J. M. Burkholder. 2013. Misuse of the phytoplankton–zooplankton dichotomy: the need to assign organisms as mixotrophs within plankton functional types. Journal of plankton research 35:3–11.

Früh, D., H. Norf, and M. Weitere. 2011. Response of biofilm-dwelling ciliate communities to enrichment with algae. Aquatic Microbial Ecology 63:299–309.

Garstecki, T., and S. A. Wickham. 2001. Effects of resuspension and mixing on population dynamics and trophic interactions in a model benthic microbial food web. Aquatic Microbial Ecology 25:281–292.

Guillard, R. R. L., and C. J. Lorenzen. 1972. Yellow-green algae with chlorophyllide c. J. Phycol. 8:10–14.

Hampl, V., I. Čepička, and M. Eliáš. 2019. Was the mitochondrion necessary to start eukaryogenesis? Trends in microbiology 27:96–104.

Hillebrand, H., C. D. Dürselen, D. Kirschtel, U. Pollingher, and T. Zohary. 1999. Biovolume calculation for pelagic and benthic microalgae. Journal of Phycology 35:403–424.

Hoshina, R., Y. Kato, S. Kamako, and N. Imamura. 2005. Genetic evidence of “American” and “European” type symbiotic algae of Paramecium bursaria Ehrenberg. Plant Biology 7:526–532.

Hsu, V., and H. V. Moeller. 2021. Metabolic Symbiosis Facilitates Species Coexistence and Generates Light-Dependent Priority Effects. Frontiers in Ecology and Evolution 8:614367.

Hsu, V., F. Pfab, and H. V. Moeller. 2022. Niche expansion via acquired metabolism facilitates competitive dominance in planktonic communities. Ecology n/a:e3693.

Hughes, E. A., M. Maselli, H. Sørensen, and P. J. Hansen. 2021. Metabolic Reliance on Photosynthesis Depends on Both Irradiance and Prey Availability in the Mixotrophic Ciliate, Strombidium cf. basimorphum. Frontiers in Microbiology 12.

Iwai, S., K. Fujita, Y. Takanishi, and K. Fukushi. 2019. Photosynthetic endosymbionts benefit from host’s phagotrophy, including predation on potential competitors. Current Biology 29:3114–3119. e3113.

Jansson, M., P. Blomqvist, A. Jonsson, and A. K. BergstrÖm. 1996. Nutrient limitation of bacterioplankton, autotrophic and mixotrophic phytoplankton, and heterotrophic nanoflagellates in Lake Örträsket. Limnology and Oceanography 41:1552–1559.

Jones, R. I. 1994. Mixotrophy in planktonic protists as a spectrum of nutritional strategies. Marine microbial food webs 8:87–96.

Kies, L. 1967. Oogamie bei *Eremosphaera viridis* De Bary. Flora B 157:1–12.

Kuznetsova, A., P. B. Brockhoff, and R. H. B. Christensen. 2017. lmerTest Package: Tests in Linear Mixed Effects Models. Journal of Statistical Software 82:1–26.

Lawrence, J., and R. Snyder. 2011. Feeding behaviour and grazing impacts of a Euplotes sp. on attached bacteria. Canadian Journal of Microbiology 44:623–629.

Laybourn-Parry, J., S. J. Perriss, G. G. R. Seaton, and J. Rohozinski. 1997. A mixotrophic ciliate as a major contributor to plankton photosynthesis in Australian lakes. Limnology and Oceanography 42:1463–1467.

Leles, S. G., L. Polimene, J. Bruggeman, J. Blackford, S. Ciavatta, A. Mitra, and K. J. Flynn. 2018. Modelling mixotrophic functional diversity and implications for ecosystem function. Journal of Plankton Research 40:627–642.

Lowe, C. D., E. J. Minter, D. D. Cameron, and M. A. Brockhurst. 2016. Shining a light on exploitative host control in a photosynthetic endosymbiosis. Current Biology 26:207–211.

Lund, J. W. G., C. Kipling, and E. D. Le Cren. 1958. The inverted microscope method of estimating algal numbers and the statistical basis of estimations by counting. Hydrobiologia 11:143–170.

Madoni, P., T. Berman, O. Hadas, and R. Pinkas. 1990. Food selection and growth of the planktonic ciliate Coleps hirtus isolated from a monomictic subtropical lake. Journal of Plankton Research 12:735–741.

Maselli, M., A. Altenburger, D. K. Stoecker, and P. J. Hansen. 2020. Ecophysiological traits of mixotrophic Strombidium spp. Journal of Plankton Research 42:485–496.

Mcmanus, G., D. Schoener, and K. Haberlandt. 2012. Chloroplast symbiosis in a marine ciliate: ecophysiology and the risks and rewards of hosting foreign organelles. Frontiers in Microbiology 3.

McManus, G. B., W. Liu, R. A. Cole, D. Biemesderfer, and J. L. Mydosh. 2018. Strombidium rassoulzadegani: a model species for chloroplast retention in Oligotrich ciliates. Frontiers in Marine Science 5:205.

Mitra, A., K. J. Flynn, J. M. Burkholder, T. Berge, A. Calbet, J. A. Raven, E. Granéli, P. M. Glibert, P. J. Hansen, and D. K. Stoecker. 2014. The role of mixotrophic protists in the biological carbon pump. Biogeosciences 11:995–1005.

Mitra, A., K. J. Flynn, U. Tillmann, J. A. Raven, D. Caron, D. K. Stoecker, F. Not, P. J. Hansen, G. Hallegraeff, and R. Sanders. 2016. Defining planktonic protist functional groups on mechanisms for energy and nutrient acquisition: incorporation of diverse mixotrophic strategies. Protist 167:106–120.

Miura, T., H. Moriya, and S. Iwai. 2017. Assessing phagotrophy in the mixotrophic ciliate Paramecium bursaria using GFP-expressing yeast cells. FEMS microbiology letters 364.

Moorthi, S. D., J. A. Schmitt, A. Ryabov, I. Tsakalakis, B. Blasius, L. Prelle, M. Tiedemann, and D. Hodapp. 2016. Unifying ecological stoichiometry and metabolic theory to predict production and trophic transfer in a marine planktonic food web. Philosophical Transactions of the Royal Society of London B: Biological Sciences 371.

Ning, Y. Z., H. F. Du, T. Zou, H. J. Wang, X. J. Wang, H. C. Liu, Z. X. Ma, and L. Ding. 2011. Allelopathy of diterpenoids on three species of soil ciliates. Acta Ecologica Sinica 31:317–321.

Pålsson, C., and C. Daniel. 2004. Effects of prey abundance and light intensity on nutrition of a mixotrophic flagellate and its competitive relationship with an obligate heterotroph. Aquatic Microbial Ecology 36:247–256.

Pröschold, T., T. Darienko, P. C. Silva, W. Reisser, and L. Krienitz. 2011. The systematics of Zoochlorella revisited employing an integrative approach. Environmental Microbiology 13:350–364.

Pröschold, T., D. Rieser, T. Darienko, L. Nachbaur, B. Kammerlander, K. Qian, G. Pitsch, E. P. Bruni, Z. Qu, D. Forster, C. Rad-Menendez, T. Posch, T. Stoeck, and B. Sonntag. 2021. An integrative approach sheds new light onto the systematics and ecology of the widespread ciliate genus Coleps (Ciliophora, Prostomatea). Scientific reports 11:5916.

Ptacnik, R., A. Gomes, S.-J. Royer, S. A. Berger, A. Calbet, J. C. Nejstgaard, J. M. Gasol, S. Isari, S. D. Moorthi, and R. Ptacnikova. 2016. A light-induced shortcut in the planktonic microbial loop. Scientific reports 6:29286.

Reisser, W. 1992. Endosymbiotic associations of algae with freshwater Protozoa and invertebrates.Plants, Animals, Fungi, Viruses, Interactions Explored. Xiii 746p. Biopress Ltd: Bristol, England, Uk. Illus. 741–719.

Rothhaupt, K. O. 1988. Mechanistic resource competition theory applied to laboratory experiments with zooplankton. Nature 333:660–662.

Rothhaupt, K. O. 1996. Laboratorary Experiments with a Mixotrophic Chrysophyte and Obligately Phagotrophic and Photographic Competitors. Ecology 77:716–724.

Rottberger, J., A. Gruber, J. Boenigk, and P. G. Kroth. 2013. Influence of nutrients and light on autotrophic, mixotrophic and heterotrophic freshwater chrysophytes. Aquatic Microbial Ecology 71:179–191.

Sanders, R. W., K. G. Porter, and D. A. Caron. 1990. Relationship between phototrophy and phagotrophy in the mixotrophic chrysophyte Poterioochromonas malhamensis. Microbial Ecology 19:97–109.

Savage, V. M., J. F. Gillooly, J. H. Brown, G. B. West, and E. L. Charnov. 2004. Effects of body size and temperature on population growth. The American Naturalist 163:429–441.

Schoener, D. M., and G. B. McManus. 2017. Growth, grazing, and inorganic C and N uptake in a mixotrophic and a heterotrophic ciliate. Journal of Plankton Research 39:379–391.

Selosse, M. A., M. Charpin, and F. Not. 2017. Mixotrophy everywhere on land and in water: the grand écart hypothesis. Ecology Letters 20:246–263.

Smalley, G. W., D. W. Coats, and D. K. Stoecker. 2003. Feeding in the mixotrophic dinoflagellate Ceratium furca is influenced by intracellular nutrient concentrations. Marine Ecology-Progress Series 262:137–151.

Stoecker, D. K., P. J. Hansen, D. A. Caron, and A. Mitra. 2017. Mixotrophy in the marine plankton. Annual Review of Marine Science 9:311–335.

Stoecker, D. K., M. D. Johnson, C. de Vargas, and F. Not. 2009. Acquired phototrophy in aquatic protists. Aquatic Microbial Ecology 57:279–310.

Team, R. D. C. 2020. R: a language and environmentfor statistical computing. R Foundation for Statistical Computing, Vienna.

Tittel, J., V. Bissinger, B. Zippel, U. Gaedke, E. Bell, A. Lorke, and N. Kamjunke. 2003. Mixotrophs combine resource use to outcompete specialists: Implications for aquatic food webs. PNAS 100:12776–12781.

Unrein, F., J. M. Gasol, F. Not, I. Forn, and R. Massana. 2014. Mixotrophic haptophytes are key bacterial grazers in oligotrophic coastal waters. The ISME Journal 8:164–176.

Urabe, J., T. B. Gurung, T. Yoshida, T. Sekino, and M. Nakanishi. 2000. Diel changes in phagotrophy by Cryptomonas in Lake Biwa. Limnology and Oceanography 45:1558–1563.

Weisse, T. 2017. Functional diversity of aquatic ciliates. European Journal of Protistology 61:331–358.

Wilken, S., J. Huisman, S. Naus-Wiezer, and E. Van Donk. 2013. Mixotrophic organisms become more heterotrophic with rising temperature. Ecology Letters 16:225–233.

Zubkov, M. V., and G. A. Tarran. 2008. High bacterivory by the smallest phytoplankton in the North Atlantic Ocean. Nature 455:224–226.

